# Evidence of spatial periodic firing in the subiculum of mice

**DOI:** 10.1101/2024.05.10.593498

**Authors:** P. Abad-Perez, G. Rigamonti, F.J. Molina-Paya, G. Cabral-Pereira, M. Esteve, R. Scott, V. Borrell, L. Martínez-Otero, A. Falco, J.R. Brotons-Mas

**Affiliations:** Universidad Cardenal Herrera-CEU, CEU Universities, Spain; Instituto de Neurociencias UMH-CSIC. Alicante; Grupo de Investigación Multidisciplinar en la enfermedad de Alzheimer UMH-UCHCEU; Universidad de Alicante, Alicante

**Keywords:** Subiculum, spatial coding, grid-like firing, place cells, boundary-vector cells, corner cells, bursting, proximodistal gradient, silicon probes, tetrodes

## Abstract

The subiculum is a critical node of the hippocampal formation, integrating multiple circuits—including thalamic inputs and afferents from CA1 and medial entorhinal cortex—and projecting broadly to cortical and subcortical targets. Yet its contribution to spatial coding remains incompletely understood. We recorded single units in freely moving mice using two complementary approaches: (i) chronic tetrodes targeting CA1 and dorsal SUB, and (ii) 64-channel linear silicon probes targeting dorsal SUB. In addition to place cells, boundary-vector cells (BVC) and corner cells (CC), we identified a subset of SUB neurons that exhibited spatially periodic (grid-like) firing. This phenomenon was replicated across recording technologies indicating that periodic coding is a consistent feature of mouse subiculum. Compared with CA1 place cells, SUB spatial neurons showed lower spatial information and reduced within-session stability, suggesting distinct coding regimes across these hippocampal subregions. Sampling along the proximodistal axis with probe arrays further revealed that burst propensity correlated positively with spatial information at more distal recording sites, consistent with known physiological gradients in subiculum and echoing relationships seen in CA1. Together, these results expand the repertoire of identified spatial codes in SUB and support a view in which subiculum contributes to geometry- and periodicity-based representations that complement CA1 and entorhinal spatial representation, thus, shaping downstream computations in cortico-subcortical circuits.

**Significance Statement:** Spatial information and memory emerge from interactions among hippocampal and entorhinal circuits with diverse, spatially tuned neurons. Here we provide the first evidence in mice that pyramidal neurons in the subiculum exhibit grid-like, spatially periodic firing, replicated across tetrodes and high-density probes. These findings suggest that the subiculum contributes to computations beyond simple relay/integration of CA1 inputs, adding a periodic component to subicular spatial coding that may shape downstream cortico-subcortical circuits.

## Introduction

Since the discovery of hippocampal place cells, neurons that code for space by being active in specific locations of the environment (1), multiple efforts have aimed to understand the computational bases of spatial navigation. As a result, different types of spatially modulated neurons have been described across brain regions: head direction cells (HD); border and boundary vector cells (BVC), coding for the geometry of the environment in medial entorhinal cortex (MEC) and the SUB; and grid cells (GC) in the medial entorhinal cortex (MEC) (2–8). These neurons are either strongly involved in the codification of external information, such as BVC coding for geometry, or in egocentric related information. This is the case for grid cells (GC), where tessellating regular firing is associated with the representation of spatial metrics and is therefore essential for path integration—the ability to navigate in the absence of external cues (9). The integration of both allocentric and egocentric streams of information is instrumental in generating an efficient spatial representation.

Multiple computational models have sought to uncover the mechanisms underlying the integration of different types of spatial signals. The BVC model suggested that place cells require the existence of neurons coding for the geometry of the environment (10). Other models suggested that place cells require GC activity (2,11). However, single unit recordings during neurodevelopment demonstrated that HD, place cells and boundary neurons appear before GC during neurodevelopment (12,13). Thus, the dependence of place cells activity on grid cell firing seems to be unclear (14).

Further research has considered the heterogeneity among different spatial cell types and their distributed anatomical localizations. In this context, the SUB may serve as a critical anatomical hub for spatial navigation (15,16). Strategically located between the EC and CA1, the SUB supports spatial information processing in a complex and heterogeneous manner. While spatial neurons in CA1 typically exhibit spatial reliability, the behavioral correlates of SUB neuron activity include location, speed, direction, and the geometry of the spatial context, while also providing a reliable representation of space across light-dark transitions (17–20). The SUB contains neurons with a diverse electrophysiological characteristic that might have different roles in memory processing (21,22).

Previous work in rats indicated the existence of grid-like, spatially tuned activity in the SUB, attributed to axonal inputs from the MEC (8). This raised the possibility that SUB pyramidal neurons could integrate periodic signals from the EC and aperiodic signals from CA1. However, whether such grid-like activity originates from local SUB pyramidal neurons has not been extensively investigated until now. Here, we sought to test the presence of periodic firing in individual SUB cells of mice using large open-field arenas. We found evidence of grid-like and other forms of spatially periodic firing (i.e., band-like,(23) in SUB neurons whose electrophysiological characteristics are compatible with pyramidal neurons. To the best of our knowledge, this is the first evidence of spatially periodic neurons in the mice SUB, highlighting the potential role of this structure in shaping the spatial cognitive map.

## Methods

### Subjects

Male wild-type C57BL/6 mice, N=11, age p60 to p90 supplied by the UMH “Servicio de Experimentación Animal (SEA)" were used. Mice were maintained on a 12 h light/dark cycle with food and water available *ad libitum* and were individually housed after electrode implantation. All procedures were approved by the UMH-CSIC ethics committee, and the regional government, and they complied with local and European guidelines for animal experimentation (86/609/EEC).

### In vivo recordings on freely moving mice

Custom-designed microdrives (Axona Ltd.) carrying twisted-wire tetrodes (12 µm tungsten; California Fine Wire, Grover Beach, CA, USA) targeted dorsal CA1 and dorsal subiculum. Under isoflurane anesthesia (1.5%) with peri-operative buprenorphine (0.05 mg/kg, s.c.), small craniotomies were made over dorsal hippocampus and dorsal subiculum. Guide cannulas were aimed at dorsal CA1 (AP −2.2 mm, ML +1.0 mm from bregma) and predominantly proximal dorsal subiculum (AP −2.7 mm, ML +0.5 mm; right hemisphere; Fig. 1A). Of the animals implanted with microdrives, three met anatomical/physiological criteria for inclusion in both CA1 and SUB datasets. Because tetrodes advance along a single guide, coverage along the proximodistal axis was limited. To increase yield and proximodistal coverage, and the number or recorded neurons, a separate cohort of two mice was implanted with 64-channel linear silicon probes (NeuroNexus, Ann Arbor, MI, USA) targeting dorsal subiculum (AP −3.8 mm, ML +1.5 mm), coordinates chosen to maximize subicular coverage. The linear arrays (≈800 µm span) enabled sampling from proximal to more distal positions within dorsal subiculum. Electrode depths (tetrodes and probes) were adjusted across days until characteristic hippocampal/subicular activity was observed.

**Figure 1.**
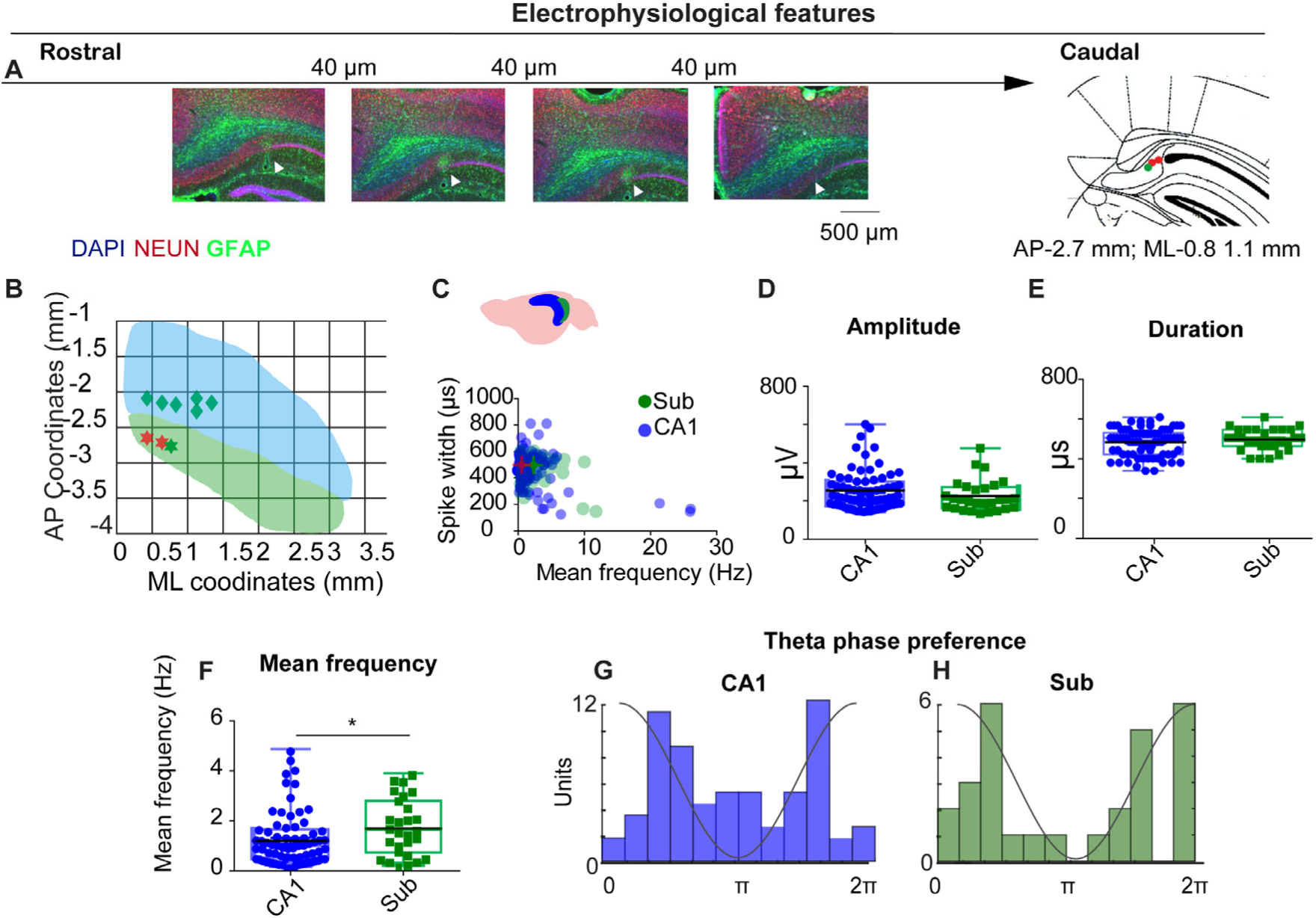
Electrophysiological features of CA1 and subiculum recordings. (A) Histological verification of tetrode tracks across the rostro-caudal extent of dorsal subiculum (DAPI/NEUN/GFAP). Arrowheads, electrode tip; schematic at right shows example target (AP −2.7 mm; ML +0.8–1.1 mm). Scale bars as indicated. (B) Recording coverage in mediolateral (ML) and anteroposterior (AP) coordinates; green = SUB, blue = CA1. (C) Scatter of spike width (trough-to-peak, µs) vs mean firing rate (Hz) for CA1 (blue) and SUB (green). (D–F) Group comparisons for spike amplitude (D), spike duration (E) and mean firing rate (F). SUB neurons fired faster than CA1 (see Results). (G–H) Theta phase preference histograms (4–12 Hz) for CA1 (G) and SUB (H); CA1 shows bimodality with a concentration near trough; SUB preferentially fires near the theta peak.

### Data acquisition

#### Tetrode recordings

As in our prior work (24) recordings were obtained using 16-channel headstage (gain x1), with an infrared light-emitting diode (LED) to track mouse position (Axona Ltd, UK). Signals were amplified (400 to 1000x), band pass filtered 0.3 Hz to 24 KHz, (Dacq system, Axona, UK) and recorded at 48 KHz/24 bit precision.. For single unit recordings, the signal was band-pass filtered between 380 Hz and 6 KHz. Spikes were recorded whenever the signal was 3-4 times above background noise, typically 20-30 µV, and stored in windows of 1 ms duration (200 ms before threshold and 800 ms after threshold detection). Spike waveforms were sampled at 48 KHz while the same channels were also recorded in continuous mode at 4.8 KHz and band-pass filtered between 0-3-2.4 KHz.

#### Silicon-probe recordings

In line with our earlier work (25), high-density recordings were obtained with an Intan-based system (NeuroNexus, Ann Arbor, MI, USA) at 30 kHz / 16-bit. The wideband stream was used for spike sorting; a downsampled 1.25 kHz copy served as the LFP signal. During sessions, electrode advancement and target verification relied on electrophysiological landmarks—ripple power, unit activity, and theta power—to optimize depth and confirm subicular/hippocampal placement.

### Place cells recordings

Mice performed a pellet-chasing task. Animals were maintained under mild food restriction (≤20% below free-feeding weight). During recording, small food pellets were tossed pseudo-randomly into the arena at ∼20s intervals to sustain locomotion and promote uniform spatial sampling. All mice were habituated to each arena for several days before recording. Tetrode-implanted mice were recorded in a 50 × 50 cm arena (20 min) and in a larger 70 × 70 cm arena (30 min). The silicon-probe cohort was recorded in an 80 × 80 cm arena (30–40 min) to maximize the chances of detecting grid/periodic firing.

### Data Analysis

Local field potential data was analyzed using custom-written MatLab codes (Mathworks, Natick, MA, USA). Raw recordings were FIR filtered (<1.2 KHz) and down sample to 2.4 KHz. Similar to previous studies (24,26), data obtained from open-field recordings was used to characterize the LFP. To visualize the power spectrum in relationship to the speed of movement, the spectral power (in decibels, 10log10) and spectrogram was built using the Thomson multi-taper method. A sliding window with 50% overlap, yielding a frequency resolution of 1 Hz was used.

### Unit isolation and spike train analysis

Single-unit activity from tetrode recordings was isolated with TINT (Axona, St. Albans, UK). This software is specifically designed for tetrode data cluster sorting. Feature vectors included the first principal components, peak-to-trough duration, peak and trough amplitudes, time of peak and trough, and waveform energy on each wire. KlustaKwik was used for initial automated clustering, followed by manual curation in TINT based on principal components, spike amplitude, and waveform consistency. Cluster quality was quantified by the overlap probability metric as described previously (24,27). Spike trains were analyzed by generating interval time histograms and temporal auto-correlogram. Only units with no spikes in the refractory period of the inter-spike time histogram (1-2 ms), and with spike amplitudes 3-4 times above background noise, typically 20-30 mV, were included. Putative pyramidal cells and interneurons were distinguished based on waveform duration and firing rate characteristics, as previously described (8). Further spike train analysis was performed to determine the bursting properties of the recorded neurons using the burst index (BI) proposed by Royer and the propensity to burst (PrtB) proposed by Staff (28,29).

For high-density recordings with silicone probes we followed the same approach as in our previous work (25). In brief, spike sorting was performed semi-automatically with KiloSort (https://github.com/cortex-lab/KiloSort) via the KilosortWrapper pipeline (https://github.com/brendonw1/KilosortWrapper), as in (9, 10, 56). Resulting clusters were manually refined in Phy (https://github.com/kwikteam/phy) using community plug-ins (https://github.com/petersenpeter/phy2-plugins). The following parameters were used for KiloSort: pos.Nfilt: 6 *numberChannels; ops.nt0: 64; ops.whitening: ‘fulll’; ops.nSkipCov: 1; ops.whiteningRange: 64; ops.criterionNoiseChannels: 0.00001; ops.Nrank: 3; ops.nfullpasses: 6; ops.maxFR: 20000; ops.fshigh:300; ops.ntbuff: 64; ops.scaleproc: 200; ops.Th: [4 10 10]; ops.lam: [5 20 20]; ops.nannealpasses: 4; ops.momentum: 1./[20 800]; ops.shuffleclusters: 1. Electrophysiological neuron classification was performed using CellExplorer (https://cellexplorer.org) (Petersen et al., 2021). This algorithm classifies units using two main features: spike waveform width (trough-to-peak, TTP) and burstiness derived from the autocorrelogram (ACG). Using the waveform and burstiness criteria, units are tentatively segregated to narrow waveform (trough-to-peak ≤ 450 μs), wide waveform (trough-to-peak > 450 μs and τ_rise_ > 6 ms), putative interneurons, and the rest, pyramidal neurons or unclassified. Because tetrode spikes were extracted from thresholded high-pass snippets whereas probe spikes were extracted from broadband continuous data, we did not compare absolute waveform amplitudes across technologies. All downstream spatial metrics (rate-map construction, information, coherence, stability) and grid/periodicity analyses were identical for tetrode and probe datasets.

### Phase locking to ongoing oscillations

Phase locking of single units to ongoing oscillations was computed as previously described (25,30). The analysis toolbox is available online (https://github.com/valegarman/HippoCookBook). The LFP was filtered for each frequency of interest, and the Hilbert transform was applied to obtain the instantaneous phase. Phase locking was computed by calculating the phase angles for each timestamp where an action potential occurred. A histogram of phase angles was calculated, and the circular mean and resultant vector were calculated for each neuron (31). Only neurons that were significantly modulated (Rayleigh test) for each frequency band during the baseline condition were included for the analysis.

#### Firing rate maps

In the case of the tetrode recordings, firing-rate maps were assembled as described previously (8,32). Pixel maps were converted to a bin matrix with a bin size of 2.5 cm x 2.5 cm. Firing rate in each bin was determined by a smoothing process using overlapping squares of 7.5 cm x 7.5 cm, as described before. Firing fields were plotted as a contour map reflecting the frequency of firing; colors were interpolated from the top firing bins down to the lowest firing areas by scaling them in decreasing intervals of the peak firing and giving them a color scale: red highest frequency, dark blue, lowest frequency. A similar approach was implemented with multisite experiments, however, in this case the FMA toolbox was used (https://fmatoolbox.sourceforge.net/.).

For both sets of data, the spatial information content per spike and per second was quantified using the Skaggs information indexes for the smoothed map (33). In addition, spatial coherence, a spatial autocorrelation measurement, was calculated as the correlation coefficient between the firing rate of each bin and the average rate of its eight surrounding bins (34). Firing fields were defined as a group of 6 contiguous bins, where the firing frequency was above the mean firing frequency plus the standard error of the firing matrix. The maximal firing frequency of this group of bins had to be above 1 Hz. For those units displaying more than one firing field, firing field size was computed as the sum of the existing firing fields and expressed as the percentage of the arena size occupied by the firing field calculated using the smoothed firing matrix. In-field maximum frequency was computed as the maximum firing field frequency of the smoothed firing map. Mean frequency was computed as the total number of spikes divided by the total recording time to provide the average session firing rate expressed in Hz.

To classify units as having spatially related activity, a randomized distribution was calculated by circular shifting the spikes times vs the position. This strategy allowed us to generate a random distribution without affecting the temporal firing structure of neurons (13). The values were obtained from the original unsmoothed firing map while respecting the original trajectory of mice (n=1000). For a unit to be regarded as being spatially modulated, it had to display a spatial coherence above 95% of the randomized distribution. This same random distribution was used to obtain the intratrial stability index between the original and a random distribution. Thus, for units to be regarded as stable they had to be above the 95% of cross-correlation between the original and the randomized distribution.

### Spatial typology analysis

Multiple spatial response types have been reported in subiculum (8,35,36). Here we specifically tested periodic (grid-like) firing and distinguished it from boundary-vector cells (BVCs), corner cells (CCs), and place cells (PCs) using the procedures below.

#### Grid analyses and periodic firing analyses

Grid analyses were implemented following similar criteria to previous studies (2). Spatial autocorrelograms were calculated for each firing map.

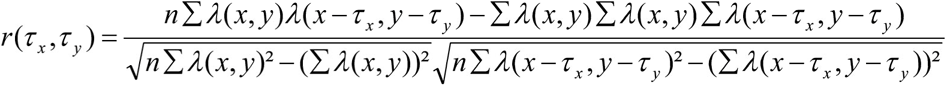

where r is the autocorrelation between the original firing rate map and its spatially shifted copy. The lag τ is normally 1-dimensional for signal analysis. Because of the 2-dimensional nature of the firing rate map, the autocorrelation has lags in the x- and y-directions: τx and τy. n is equal to the number of pixels. To determine the existence of a hexagonal grid firing structure, grid index, size and orientation were calculated as described previously (8,37).

To obtain the “gridness index” we obtained a circular portion of the spatial autocorrelogram generated from the spatial firing matrix. This circle was centered on the central peak which was eliminated from the matrix. Then we correlated the rotated versions, in steps of 6°, of this matrix with the original spatial autocorrelogram. The degree of spatial periodicity (gridness’ or ‘grid scores’) was determined for each recorded cell and the grid index calculated as previously described (37,38).

Additionally, to further investigate the presence of periodic firing we obtained the sum of values of the 2D resulting map in all its orientation and we displayed the preferred orientation of these neurons in space as a polar plot (23). These values were plotted against the polar plot of the shuffled original firing map, n=1000. Only neurons that showed higher values than this confidence interval were considered as periodic neurons.

#### Boundary vector cells (BVC) and Corner cells (CC)

To identify BVC we follow the same methodology as previously described (39). This analysis implied the comparison of the original map with idealized BVC. Spatial shuffling was performed as described above. Model BVCs were fit to these spatially shuffled rate maps as described above. To be classified as a BVC, the r_(max)_ value for each cell had to surpass the 95th percentile of the 1000 shuffled r_(max)_ values generated for that specific cell and trial while meeting the same criteria to be spatially modulated.

Corner cells were identified following the same procedure as described by Sun et al, (36). Field location was taken as the regional maximum (x,y) of each region. Centroid and corner coordinates were detected manually verified. For each field we computed a corner score

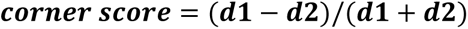

Where *d1* is the distance from the field maximum to the environment centroid and *d2* is the distance to the nearest corner. The score ranges from **−1** (centroid) to **+1** (ideal corner). The total corner score of the cell was calculated as the mean of the corner scores of the firing fields and that we considered corner cell only if the total index was above 95th of the shuffling distribution in addition the neuron had more than one firing field. Units that did not meet GC, BVC or CC were deemed as place cells (PC).

#### Histological Analysis

After completing experiments, animals were sacrificed using sodium pentobarbital (180 mg/kg at a concentration of 100 mg/ml) and transcardially perfused with saline and 4% paraformaldehyde. This procedure was authorized by the ethical commission according to the current normative and following veterinary recommendations. The brains were removed, sliced in 40 µm sections, and prepared for immunohistochemistry. For primary antibodies we used anti-NeuN (anti-NeuN 1:500; Millipore #MAB377) and GFAP (Merck, SAB2500462-100UG; host: goat; dilution 1:1000), and for secondary Anti-goat, (Invitrogen, A11055; dilution 1:1000) for electrode localization. Nuclei were stained with DAPI (Sigma-Aldrich, D9542-10ML). Electrode tracks were identified when visible and their position estimated by comparison with anatomical landmarks using the Paxinos and Franklin stereotaxic atlas. In addition, the presence of local electrophysiological signatures (e.g., ripple activity, theta oscillations) provided further confirmation of electrode placement. In those cases where a clear electrode track was not detectable, the location of the electrodes was inferred based on implant coordinates, electrode design, and characteristic electrophysiological landmarks.

#### Statistical analysis

Distribution of dependent variables was tested for normality and for equal variance. Parametric statistical tests were used for normally distributed variables and non-parametric tests were used for variables that did not show normal distribution. All descriptive values are expressed as mean ± the standard error (S.E.). Statistical significance was set at p < 0.05. All statistical analysis was performed using SPSS (IBM, V27) software package and MATLAB statistic toolbox, (Mathworks, Natick, MA, USA).

## Results

### Tetrode recordings in the CA1 and SUB

To test for spatially periodic cells in subiculum, we recorded in standard (50 × 50 cm) and large (70 × 70 cm) open fields. Across eight mice (five implanted in CA1, one in SUB, and three in both regions), histology confirmed the placement of three tetrodes in three different mice in the SUB (Fig. 1A-B) we recorded 147 CA1 and 65 SUB neurons. Units were classified by spike duration, waveform shape, and firing rate (Fig. 1C), following established criteria (7). Of these, 74 CA1 and 28 SUB cells met pyramidal and spatial-modulation criteria (Methods).

Unit isolation quality did not differ between regions (CA1 vs. SUB overlap probability: 0.06 ± 0.01 vs 0.10 ± 0.02; Mann–Whitney U = 4290, *p* = 0.236), like previous reports (24,27). Spike amplitude and duration were also similar (duration: 477.5 ± 7.8 µs vs 491.0 ± 10.4 µs, U = 862, p = 0.192; amplitude: 254.1 ± 11.9 µV vs 226.1 ± 15.6 µV, U = 943, p = 0.48; Fig. 1C–). In contrast, SUB neurons fired faster than CA1 neurons (mean rate: 1.8 ± 0.2 Hz vs 1.2 ± 0.12 Hz; U = 745, p = 0.02; Fig. 1F), consistent with prior reports (7,18,40).

The interaction between the CA1 and the SUB is critical for the organization of inputs and outputs of the hippocampal formation. This dialog depends on the specific timing of the activity of different populations of neurons in different theta cycle windows (41). Therefore, examining the relationship between CA1 and SUB firing and local LFP is fundamental to understanding the temporal dynamics of the hippocampal formation. We investigated the relationship between the firing of CA1 and SUB neurons with the locally recorded theta (4-12Hz). To target the pyramidal cell layer, we selected tetrodes on which ripples showed no slow (positive or negative) component, a standard indicator of pyramidal-layer placement. In CA1, preferred firing phases were bimodally distributed, with units clustering around the ascending and descending phases of the local theta cycle and an additional concentration near the trough (Fig. 1F). For subiculum, we applied the same tetrode-selection criterion and computed phase from the local LFP on the same tetrode; subicular neurons preferentially fired near the theta peak in the tetrode database (Fig. 1G).

### A group of subicular neurons showed grid and spatially periodic firing

In rats, subicular spatial coding comprises a heterogeneous mix of place cells (PC) and boundary-vector cells (BVCs), as well as corner cells (CC) (5,8,36). Given the larger body size of rats relative to typical arena dimensions, spatial periodicity may have been difficult to detect. In mice recorded in 50 × 50 cm arenas, we observed that several SUB neurons exhibited spatial periodic firing resembling grid-like structure. In contrast to prior reports (8), these units had pyramidal-like waveforms, were stable within session, and were well isolated (Fig. 2A–D). Their spatial firing showed hexagonal periodicity by both autocorrelogram and frequency-domain analyses (Fig. 2E–G). CA1 pyramidal neurons recorded in the same arenas did not show comparable periodic patterns (Fig. 2H–N). Overall, 4 of 28 spatially modulated SUB units (14%) met our gridness criterion (three cells in one mouse and one cell in a second mouse).

**Figure 2.**
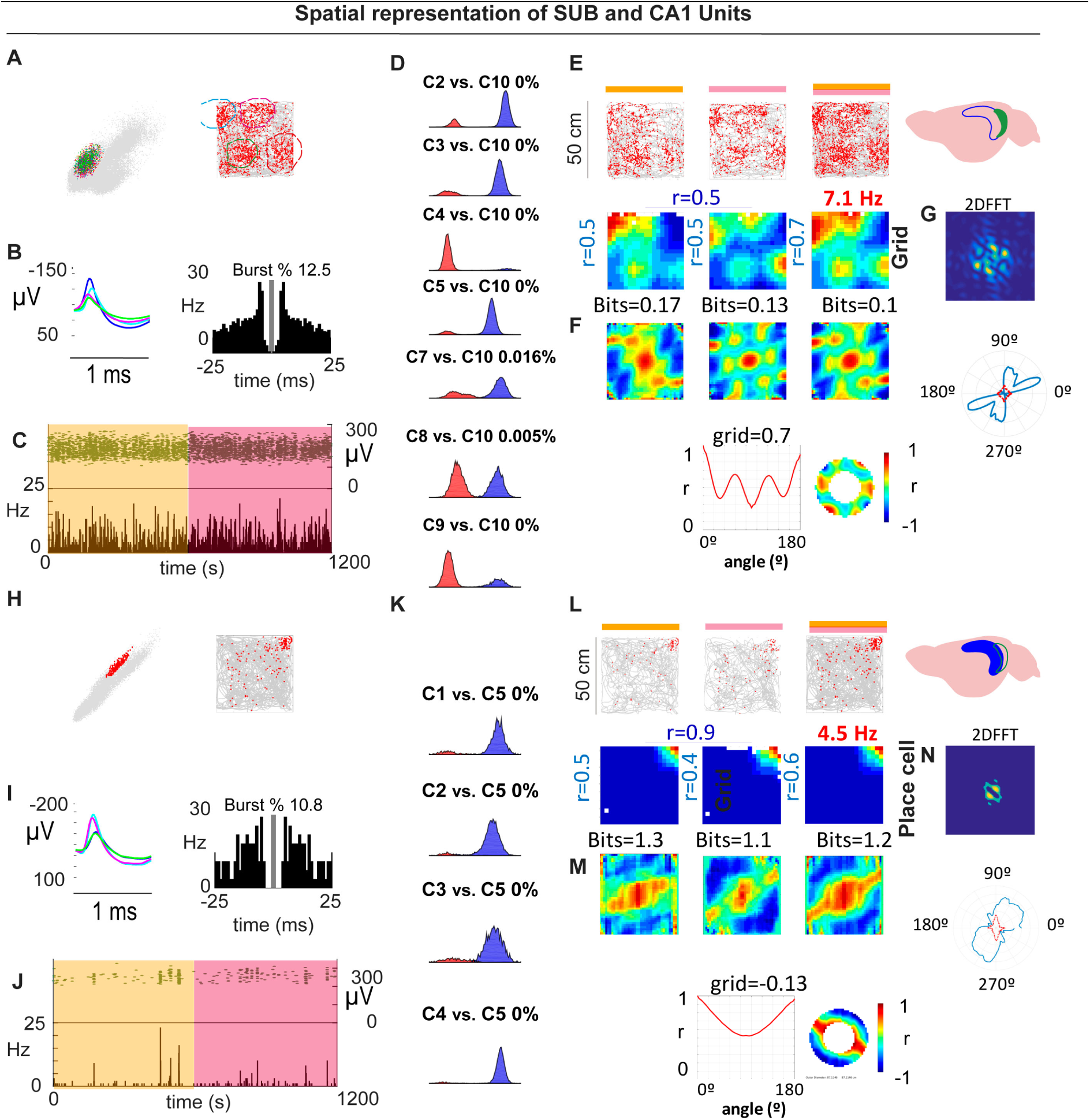
(A–C) Example SUB periodic cell: cluster isolation, waveform/ISI with burst %, and stable firing across the session.(D) Overlap probability controls for isolation quality (example comparisons). (E–F) Rate maps and autocorrelograms across spatial bins with corresponding gridness/periodicity summaries and 2D FFT (G). (H–J) Example CA1 place cell recorded in the same arena: isolation (K), rate maps/autocorrelograms (L, M) and 2D FFT (N). CA1 pyramidal units did not exhibit periodic/grid signatures under these conditions.

To test the robustness of subicular periodicity across scales, we tracked individual SUB neurons across 50 × 50 cm and 70 × 70 cm boxes. Some cells deviated from perfect hexagonal symmetry in the larger arena yet retained clear periodic structure, similar to observations in MEC under certain conditions (23). Fig. 3A–E vs. 3A′–E′). Conversely, we identified at least one cell that showed a single dominant place field in the 50 × 50 cm arena but developed periodic firing in the 70 × 70 cm arena, as revealed by the 2D FFT and related metrics (Fig. 3F–J vs. 3F′–G′).

**Figure 3.**
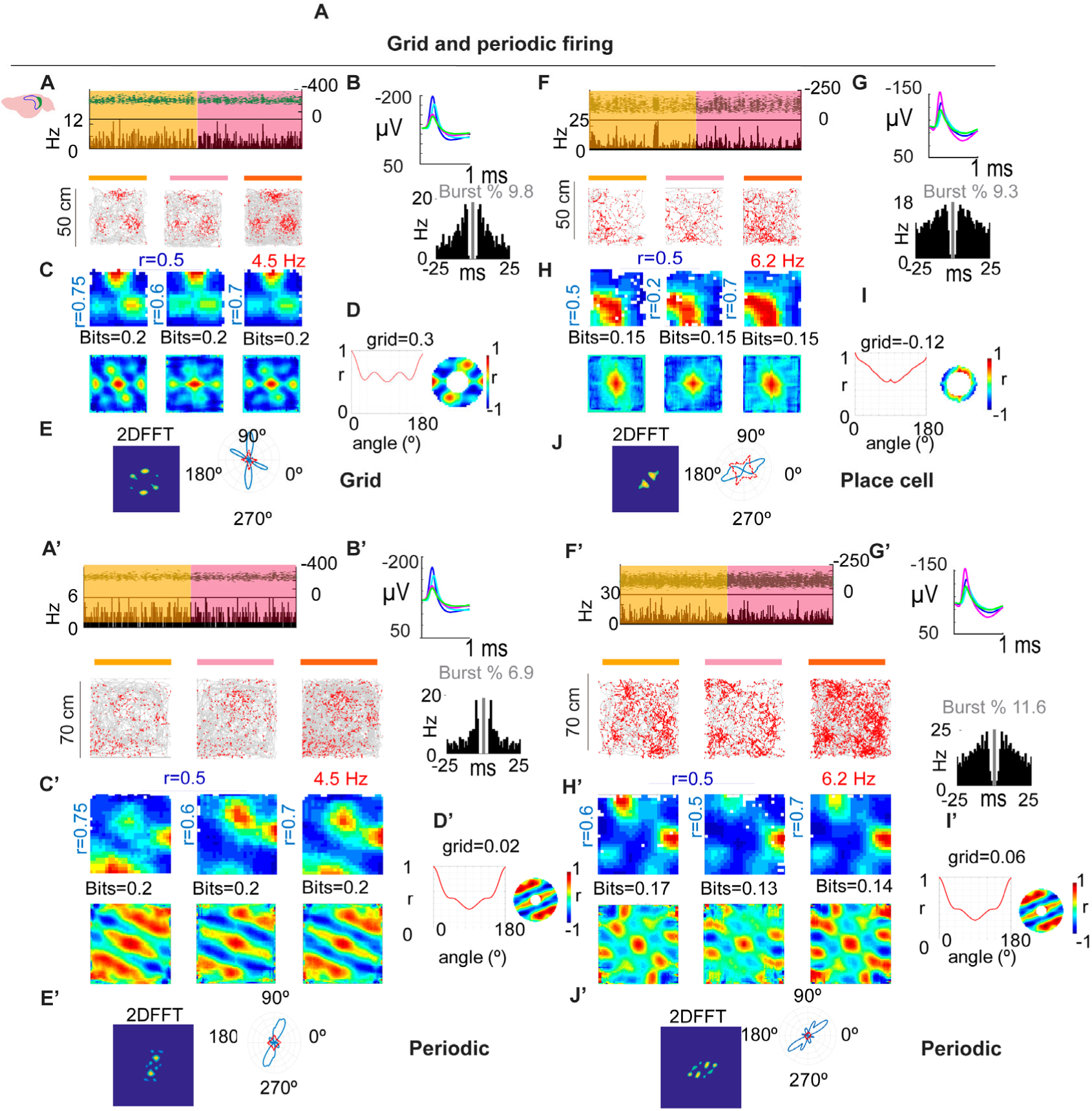
Grid and spatially periodic firing across arena size. (A–D) Example SUB neuron in 50 × 50 cm: stable firing (A), waveforms/ISI (B), rate maps and autocorrelograms (C), grid/periodicity summary (D) with 2D FFT (E). (A′–E′) Same neuron in 70 × 70 cm: periodic structure persists but deviates from perfect hexagonal symmetry (see main text). (F–J, F′–J′) Second example showing a place-like pattern in the small arena (G) that becomes periodic in the large arena, as confirmed by the 2D FFT (J, J′).

### Spatial firing of SUB neurons showed lower spatial modulation than CA1

Spatial coding differs across hippocampal subregions (19,42). We compared CA1 and SUB using standard metrics spatial information per second, spatial information per spike, spatial coherence, and firing-field size. We found significant differences between CA1 and SUB neurons for the spatial information per second, spatial information per spike, and for the spatial coherence, consistent with previous reports (Fig.4 A-C). In contrast, (Fig. 4D), no significant differences were observed for the firing field size (CA1 vs SUB : bits/s 0.39 ± 0.01 vs. 0.322 ± 0.06 µs,: U=738.0, p=0.025; bits/spike 0.43 ± 0.04 vs. 0.32 ± 0.06: U=611.0, p=0.001, Mann-Whitney; spatial coherence: (r), 0.61 ± 0.01 vs. 0.54 ± 0.02: t_(100)=_2.442, p=0.016; firing field size (%), 29.02 ± 0.7 vs. 30.0 ± 1.14, t_(100)=_-0.634, p=0.528).

**Figure 4.**
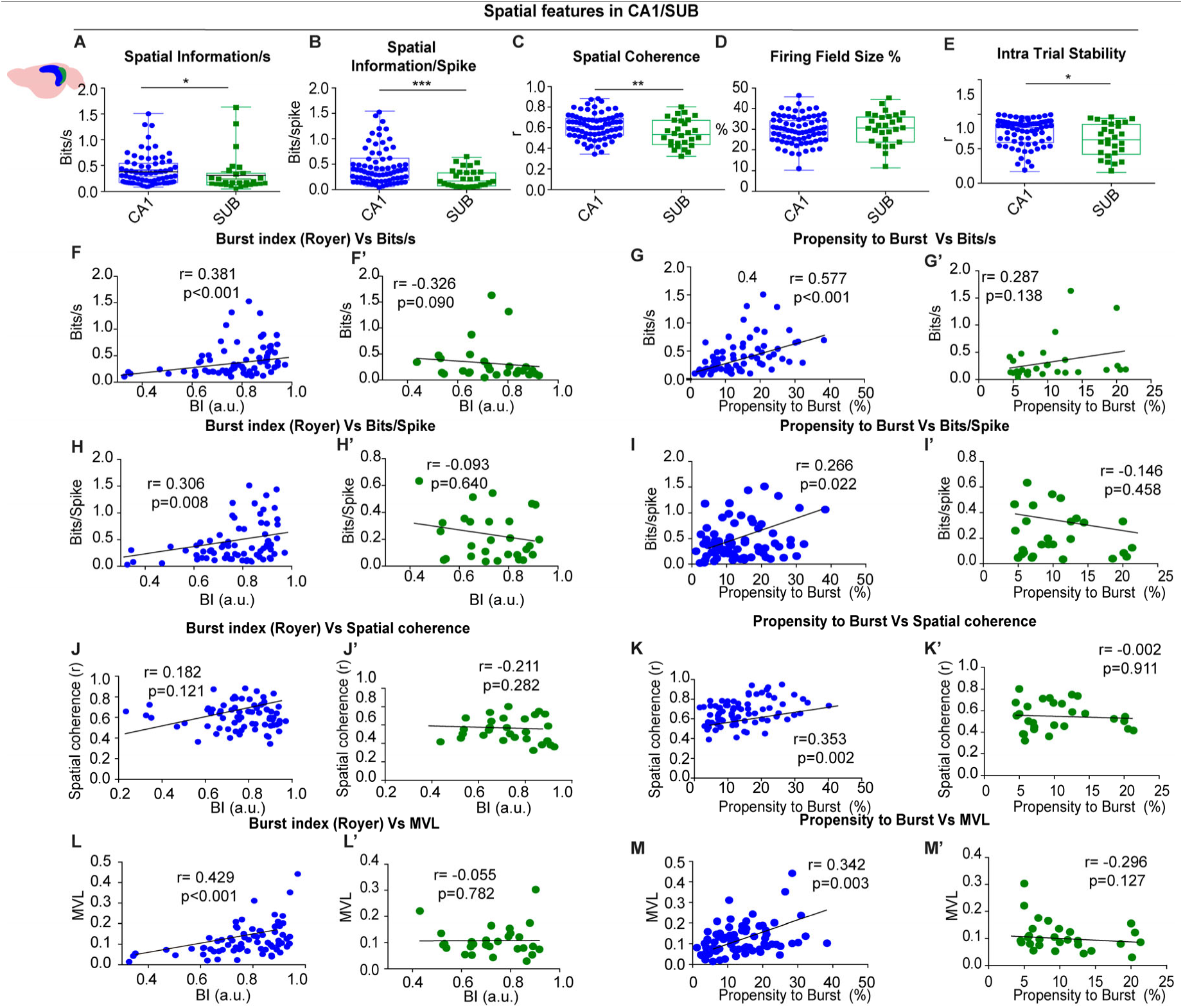
Spatial features in CA1 vs SUB (A–E) Group comparisons: spatial information per second, spatial information per spike, spatial coherence, firing-field size, and intra-trial stability and intra-trial stability were higher in CA1 (F–M′) Correlations between burst index or propensity to burst and spatial metrics (bits/s, bits/spike, coherence, MVL) for CA1 (blue) and SUB (green).

Because temporal instability can mimic rate remapping and obscure periodicity (43) we assessed within-session stability by splitting each session into two 10-min halves, (Fig 4E) and computing bin-by-bin spatial correlations. We found that spatially periodic signals that place cells in the CA1 region showed significantly higher spatial consistency (CA1 vs SUB: intra trial stability coherence (r), 0.72±0.02 vs. 0.61±0.04: U=764.0, p=0.041, Mann-Whitney). However, periodic neurons showed high stability within each trial, (See Fig 3 for different examples), ruling out remapping effects as a confounding factor in the identification of spatially-periodic firing.

### Spatial modulation of SUB units in the proximal region was not determined by bursting activity

The spatial information is closely related to the electrophysiological characteristics of hippocampal pyramidal neurons. Specifically, place cells are known to exhibit complex spikes, or bursting activity, which is characterized by the generation of trains of action potentials within an envelope of 15-20 ms. This bursting pattern is thought to enhance the fidelity of spatial information across different brain regions (44). We investigated if there was a specific relationship between bursting patterns and the spatial modulation in the CA1/SUB axis.

We found that in CA1 bursting metrics correlated with spatial modulation (Fig. 4F–M). The burst index (BI) correlated with information per second (r = 0.381, p < 0.001, Spearman) and information per spike (r = 0.306, p = 0.008), showed a trend for spatial coherence (r = 0.182, p = 0.18, Spearman), and was strongly related to theta locking (MVL; r = 0.429, p < 0.001, Spearman). The propensity to burst (PrtB) showed a similar pattern: bits/s (r = 0.577, p < 0.001), bits/spike (r = 0.266, p = 0.022, Spearman), spatial coherence (r = 0.353, p = 0.002, Spearman), and MVL (r = 0.342, p = 0.003, Spearman). These relationships indicate that stronger bursting is associated with stronger spatial coding and tighter theta–phase locking in CA1. Remarkably, we observed no correlation between bursting properties and spatial modulation in SUB-recorded units, (Fig.4F’-M’) (BI vs bits per second, r=-0.326, p=0.09; BI vs bits per spike r= -0.093, p=0.640; BI vs spatial coherence, r= -0.211, p=0.282; BI vs mean vector length, r= -0.055, p=0.782, Spearman). A similar pattern was observed when we compared the bursting index with the spatial modulation (PrtB vs bits per second, r= 0.287, p=0.138; PrtB vs bits per spike r= -0.146, p=0.458; PrtB vs spatial coherence, r= -0.002, p=0.991; PrtB vs mean vector length, r= -0.296, p=0.127, Spearman).

This indicates that CA1 neurons with a higher tendency to burst exhibited greater spatial modulation, whereas in SUB-recorded neurons, bursting activity did not predict spatial modulation, whether periodic or not.

### Multisite Recordings

To validate and extend the tetrode findings, we implanted two additional mice with 64-channel silicon probes targeting dorsal subiculum (see Fig 5A). Data were sorted with KiloSort2 and manually curated in Phy2; cell-type labels were assigned with CellExplorer (Petersen et al., 2021; see Methods). We isolated 196 with clear refractory periods: pyramidal neurons (PYR), *n* = 142; narrow-waveform interneurons (NW), *n* = 35; and wide-waveform interneurons (WW), *n* = 15, (see Fig 5B). Four units were deemed unclassified.

**Figure 5.**
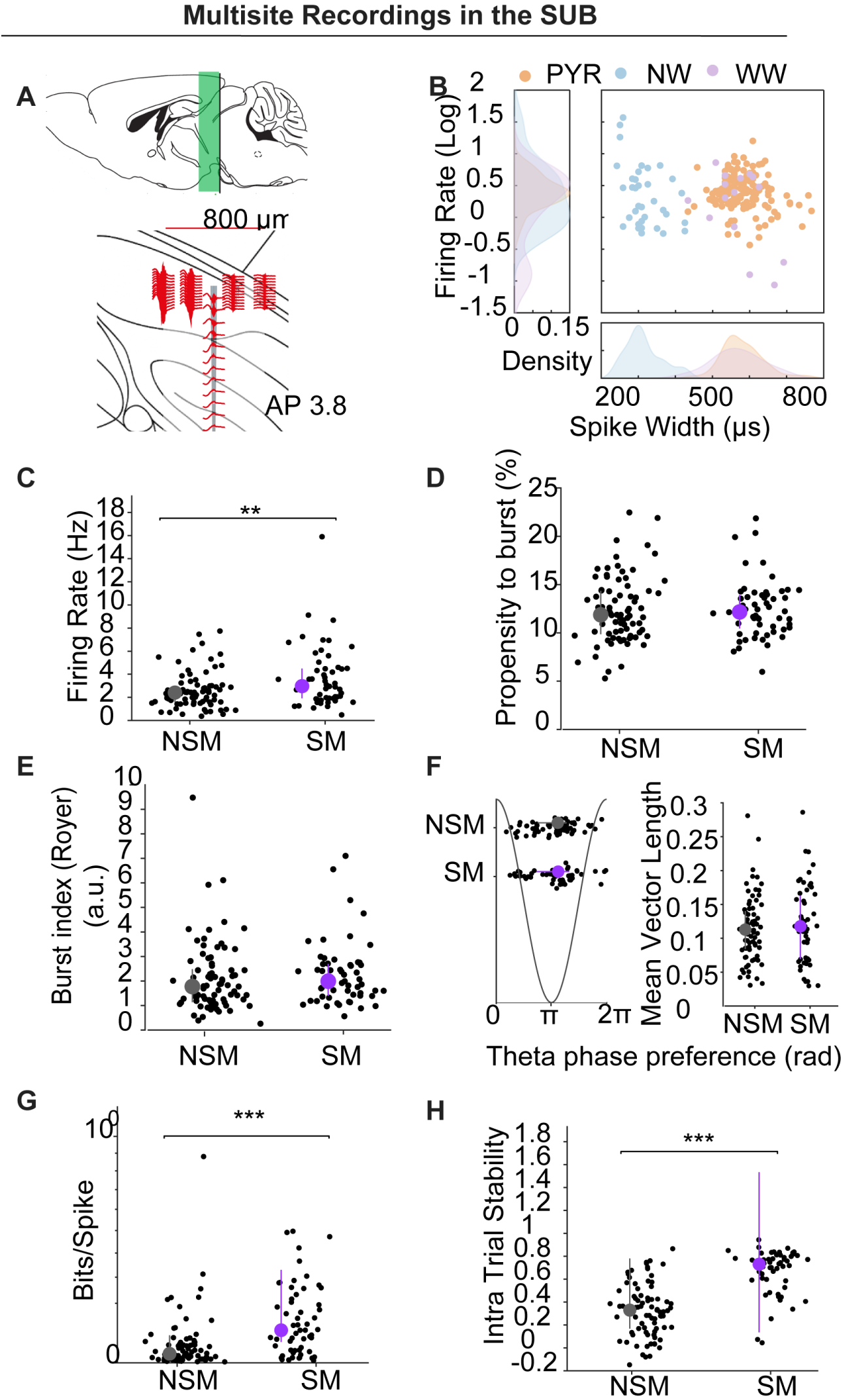
Multisite silicon-probe recordings in subiculum. (A) Schematic of probe geometry and dorsoventral span (∼800 µm) targeting dorsal SUB (AP −3.8). (B) Cell-type classification by spike width and firing rate: pyramidal (PYR, orange), narrow-waveform interneurons (NW, blue), wide-waveform interneurons (WW, purple); density plots shown at margins. (C–H) Comparisons for firing rate (C), propensity to burst (D), burst index (E), theta phase preference/locking (F), spatial information (bits/spike) (G), and intra-trial stability (H) between NSM and SM pyramidal neurons. SM fired faster, had higher bits/spike and spatial stability with no differences in PrtB, BI, or theta metrics.

### Spatial modulation

A unit was classified as spatially modulated (SM) if its rate-map spatial coherence exceeded the 95th percentile of a 1,000-iteration time-shift shuffle (See Methods). In total, 67/196 units (34.2%) met this criterion. Unless noted otherwise, the analyses below are restricted to pyramidal neurons.

Firing rate was higher in SM than in non-spatially modulated (NSM) PYR (Frequency 3.672 ± 0.347 Hz vs 2.585 ± 0.173 Hz; U = 4967, p = 0.0065; Mann–Whitney; Fig. 5C). By contrast, burst metrics—propensity to burst (PrtB) and burst index (BI)—and theta phase preference/locking did not differ between SM and NSM (PrtB: 2.441 ± 0.399 vs 12.389 ± 0.377; U = 5545, p = 0.849; Mann–Whitney Fig. 5D; BI: 2.286 ± 0.174 vs 2.081 ± 0.159; U = 5251, p = 0.139; Mann–Whitney, Fig. 5E); Theta phase (mean ± circSD, 3.443 ± 1.086 vs. 3.424 ± 0.978, Watson–Williams test: F(1) = 0.007, *p* = 0.935; Theta MVL: 0.114 ± 0.008 vs. 0.120 ± 0.006, U test: U = 4879, *p* = 0.524 Mann–Whitney, Figure F). As expected, SM units showed greater spatial information and within-session (intra-trial) stability than NSM (bits/spike: 0.145 ± 0.017 vs 0.062 ± 0.012, *U* = 4469, *p* < 0.0001; stability: 0.672 ± 0.025 vs 0.329 ± 0.025, *U* = 3877, *p* < 0.0001; Fig. 5G–H).

### Incidence of periodic cells and other spatial classes in multisite recordings

In the probe cohort, 7/67 SM units (10.4%) met our a priori grid/periodicity criteria (four and three cells in the two animals, respectively; Fig. 6A–C). Five of these were PYR and two were NW interneurons (Fig. 6C). This incidence closely matches the tetrode dataset (4/28 = 14.3%), under identical detection and shuffling thresholds, indicating that periodic firing in mouse subiculum replicates across animals and recording methods. We also identified **place cells (PC)** 36/67 (53,7%), **boundary-vector cells (BVC)** 19/67 (28.4%), and **corner cells (CC)** 5/67 (7.5%) (Fig. 6D to 6F).

**Figure 6.**
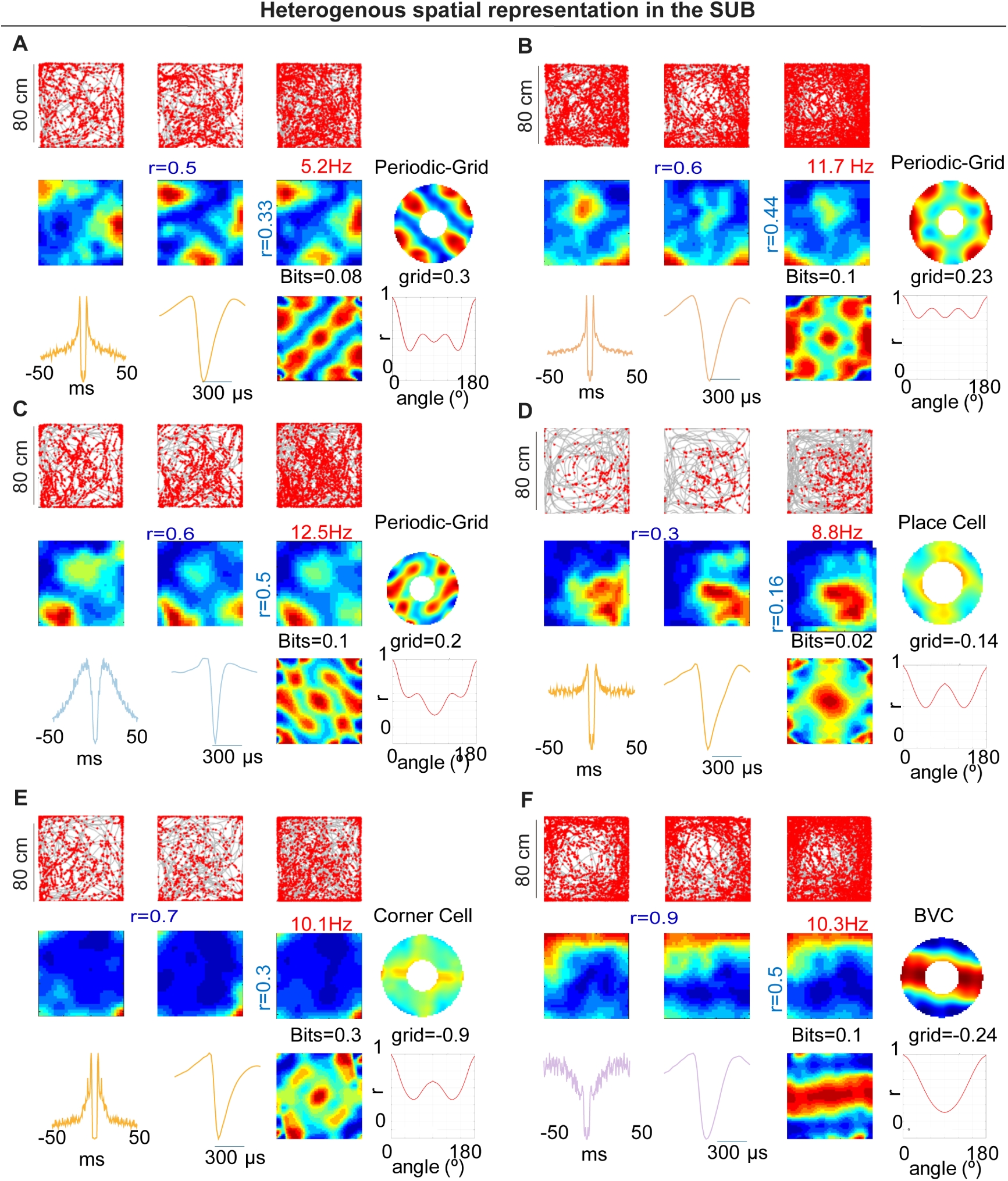
Heterogeneous spatial representations in subiculum (probe cohort, 80 × 80 cm). (A–C) Examples of periodic/grid-like neurons: trajectories with spikes, rate maps across time segments, spatial autocorrelograms, periodicity summaries (polar/orientation energy), and spike waveforms/ISI. (D) Example place cell. (E) Example corner cell with high corner-score field. (F) Example boundary-vector cell (BVC). For each neuron, grid score/periodicity measures and bits values are indicated on the panels. Across the cohort, spatial classes (PC, BVC, CC, GC, NSM) were distributed along the proximodistal axis.

We next investigated potential difference across neuron types in multiple parameters, for this analysis we included all types of neurons, with independence of its electrophysiological type (Fig 7A–D). We found that spatial information per second was higher in PC, BVC, and CC than in NSM (NSM 0.080 ± 0.011; PC 0.449 ± 0.046; BVC 0.497 ± 0.092; CC 0.322 ± 0.099; GC 0.226 ± 0.094; χ²_(4)_ = 96.523, p < 0.0001, Kruskal–Wallis, Post-hoc (Dunn–Holm): PC > NSM (adjusted p < 0.0001), BVC > NSM (adjusted p < 0.0001), CC > NSM (adjusted p = 0.035). Spatial information per spike also differed among classes: NSM 0.052 ± 0.009; PC 0.156 ± 0.021; BVC 0.121 ± 0.029; CC 0.131 ± 0.061; GC 0.047 ± 0.022; χ²_(4)_ = 42.985, p < 0.0001, Kruskal–Wallis. Post-hoc: PC > NSM (adjusted p < 0.0001), BVC > NSM (adjusted p = 0.004), PC > GC (adjusted p = 0.031). Spatial coherence was higher for all spatial classes when compare with NSM NSM 0.095 ± 0.008; PC 0.392 ± 0.021; BVC 0.441 ± 0.033; CC 0.369 ± 0.027; GC 0.308 ± 0.047; χ²(4) = 116.918, p < 0.0001; Kruskal-Wallis test. Post-hoc: PC > NSM (adjusted p < 0.0001), BVC > NSM (adjusted p < 0.0001), CC > NSM (adjusted p = 0.003), GD > NSM (adjusted p = 0.005).

**Figure 7.**
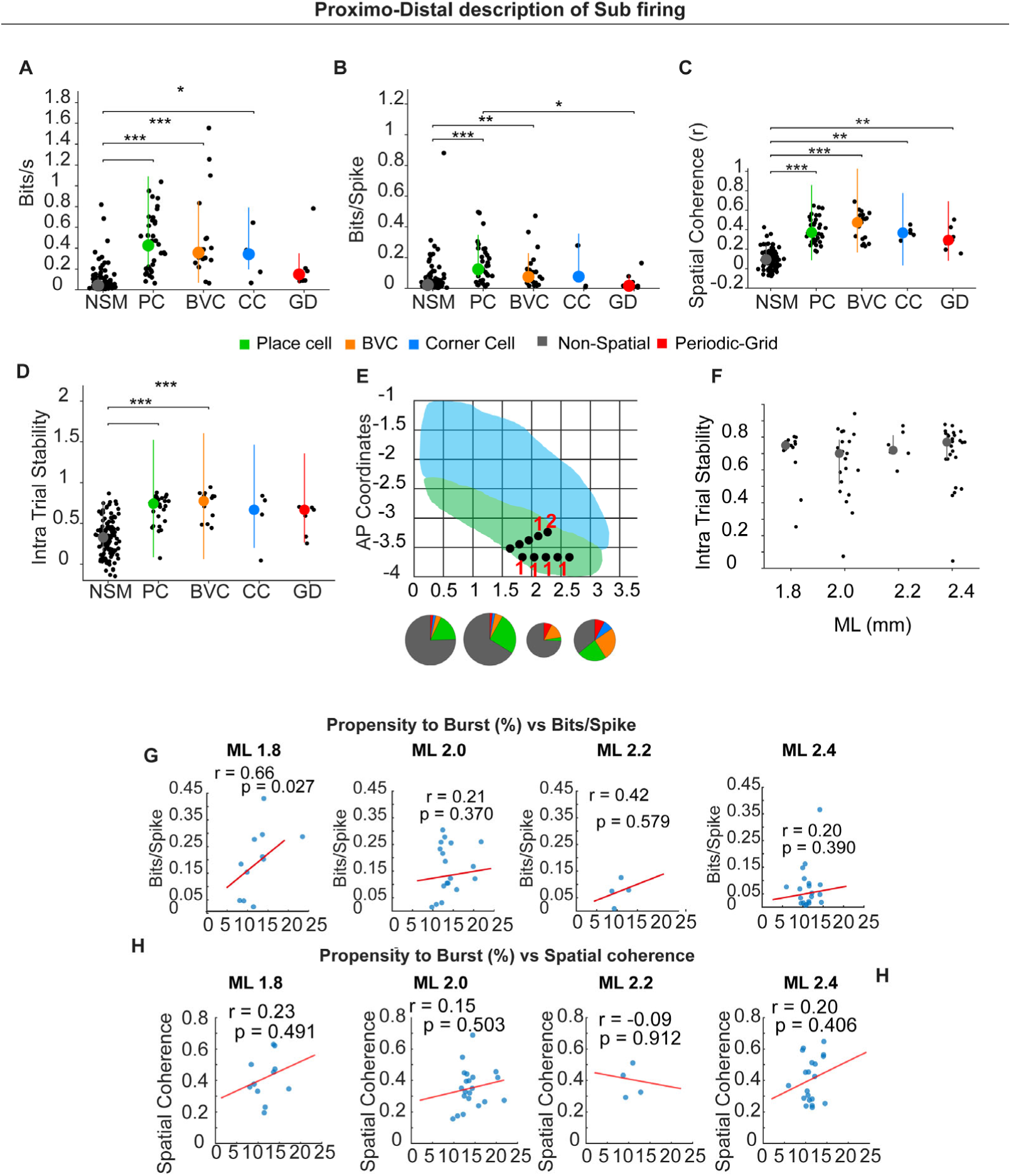
Different spatial types in the sub and Proximo–distal description of subicular firing. (A–D) Summary of spatial metrics across cell classes: NSM (non-spatial), PC (place cell), BVC (boundary-vector cell), CC (corner cell), and GD (grid/periodic). (E) Schematic of dorsal subiculum showing mediolateral (ML) and anteroposterior (AP) CA1 (blue) SUB (green). Black dots indicate recording sites; pie charts show cell-type composition within each ML bin representing numbers giving in the result text. (F). Intratrial stability across the proximo-distal axis. (G–H) Correlations (spatially modulated pyramidal neurons only) between propensity to burst (%) and spatial information (bits/spike) (F) or spatial coherence(G) computed within ML bins (1.8, 2.0, 2.2, 2.4 mm). Lines show least-squares fit; r and p are Pearson values. A significant positive relation appears distally (ML 1.8: r = 0.66, p = 0.027); other anatomical locations show no significant correlation.

We also verified that there were no significant differences in intra-trial spatial stability between spatial types. On the other hand, they presented higher spatial stability than non-spatially modulated neurons, although this effect was significant only for PC and BVC (NSM r = 0.3199 ± 0.02; PC r = 0.68 ± 0.02; BVC r = 0.74 ± 0.03; CC r = 0.58 ± 0.14; GC r = 0.56 ± 0.07; χ² _(4)_ = 84.93, *p* >0.0001, Kruskal–Wallis; Post-hoc PC > NSM (adjusted p < 0.0001), BVC > NSM (adjusted p < 0.0001); CC vs NSM (adjusted p = 0.161); GC vs NSM (adjusted p = 0.250). Spatial cell types occurred throughout the sampled proximodistal span (Fig. 7E).

### Proximodistal axis spatial stability and bursting–spatial coupling

We investigated if there were differences in the intratrial spatial stability across the proximodistal axis. For this analysis we included all spatially modulated neurons with independence of their electrophysiological profile. We found no significant differences in these parameters (1.8 mm: 0.6920 ± 0.0426; 2.0 mm: 0.6383 ± 0.0451; 2.2 mm: 0.7397 ± 0.0345; 2.4 mm: 0.6988 ± 0.0375 χ²(3) = 2.148, p = 0.542, Kruskal–Wallis). Similar results were obtained for only pyramidal neurons, (data not shown).

In the tetrode dataset—which predominantly sampled proximal dorsal subiculum—burst metrics (BI, PrtB) did not correlate with spatial information. Given the known proximal→distal gradient in bursting propensity, we re-examined this in the probe dataset, which sampled ∼800 µm along the proximodistal axis (Fig. 7G–H). Considering spatially modulated (pyramidal only), bits/spike correlated positively with burst propensity in the most distal bin (ML 1.8: r = 0.66, p = 0.027), but not in more proximal bins (ML 2.0: r = 0.21, p = 0.370; ML 2.2: r = 0.42, p = 0.579; ML 2.4: r = 0.20, p = 0.390) (Fig. 7).

## Discussion

We provide the first evidence in mice that subicular pyramidal neurons exhibit grid-like/spatially periodic firing, using two independent technologies, custom tetrodes and high-density linear silicon probes. In the probe cohort, 7/67 spatially modulated cells (10.4%) met our grid/periodicity criteria (5 pyramidal, 2 interneurons), an incidence that closely matched the tetrode dataset (4/28 = 14.3%), indicating robustness and replicability across methods. Unit isolation met established sorting-quality benchmarks (23,27,45), and isolation quality did not differ between CA1 and SUB. Finally, spike waveforms and firing statistics were consistent with pyramidal neurons and inconsistent with putative axonal recordings (46).

Multisite (silicon-probe) recordings yielded higher unit counts and broader proximodistal coverage within dorsal SUB, enabling identification of multiple spatial classes—place cells, boundary-vector cells, and corner cells—all present across the proximodistal axis, and to understand better how bursting and firing rate might contribute to spatial information transmission.

### SUB spatial representation is different to CA1

We observed that place cells in the CA1 showed higher spatial resolution as compared to SUB cells. This apparent lower spatial modulation of SUB neurons is related to their higher firing rate. Such augmented activity seems to serve as a mechanism to improve information routing from hippocampal formation to other brain regions, (47).

In the tetrode dataset, SUB neurons tended to fire near the center/peak of the theta cycle, which contrasts with prior reports showing a bias toward the descending phase and earlier than CA1 (47). This discrepancy may reflect recording location across the deep– superficial axis sampling. Consistent with this interpretation, the multisite probe recordings, which provide better laminar localization, revealed a phase preference like the published pattern (47). Thus, the apparent mismatch in the tetrode data likely stems from limited control over laminar position.

### Proximodistal organization in the Subiculum

Inputs to the SUB are topographically organized, this way the proximal region of the SUB receives information about inputs from LEC, known to be involved in the temporal coding of episodes (48), while more distal regions integrate inputs from the MEC involved in spatial coding (37), see O’ Mara et all for a comprehensive review (49,50). In addition, there is proximodistal gradient of burstiness, from lower in the proximal to higher in the distal SUB, that might indicate a segregation of functions (51).

Our multisite recordings show that the dorsal subiculum hosts a diverse ensemble of spatial codes rather than a single canonical type. Within the same animals we observed place cells, boundary-vector cells (BVCs) that anchor firing to the geometry of nearby boundaries, corner cells (CCs) with fields concentrated near vertices, and a subset of neurons with spatially periodic (grid-like) activity. Importantly, these classes were distributed along the proximodistal axis, rather than confined to a mediolateral specific coordinate, indicating that geometric and periodic representations are features of subicular circuitry. This heterogeneity suggests that SUB integrates inputs from CA1 and entorhinal cortices to form complementary codes—spatial, boundary-anchored (BVC/CC) and periodic tiling—that can support navigation and context generalization across environments.

The presence of these codes across the proximodistal extent also implies parallel output channels (49,50) from SUB to downstream cortico-subcortical targets, enabling these different signals to be differentially routed rather than relying on a single encoding scheme. This could be critical for the involvement of the sub in different cognitive processes. In this sense, SUB neurons seem capable of maintaining active information during the inter-trial in a working memory task (52). Such sustained or modulatory activity could help signal the temporal structure of episodes, with distinct proximal–distal inputs contributing to the organization of different information streams.

In our proximal SUB, tetrode recordings, spatial stability was lower than in CA1, consistent with stronger LEC influence. However, in the multisite dataset we found no significant proximodistal differences in intra-trial stability, suggesting that stability per se is relatively uniform along the axis as well as also across different spatial cell types (BVC/CC vs periodic).

It is relevant to mention that unlike CA1, where bursting often correlates with spatial information and is thought to enhance synaptic reliability, tetrode recordings in the subiculum did not show a uniform burst–spatial coupling. However, our multisite recorded data resolved this discrepancy: burst propensity correlates with spatial information only at more distal recording sites, whereas proximal SUB shows no such relationship. This proximodistal dissociation suggests two coding regimes within SUB. Proximally, spatial information is supported chiefly by elevated firing rates—a strategy that can improve signal-to-noise and downstream reliability through temporal averaging and redundant spiking. Distally, bursting contributes additional coding power beyond rate aligning distal SUB more closely with CA1-like burst–information coupling. Given the known gradients in intrinsic excitability, burst propensity, and projection targets along SUB (51), these results are consistent with a functional multiplexing: rate-dominated coding proximally and burst-augmented coding distally. Such organization could tailor information transfer to distinct downstream circuits, with bursts selectively engaging synapses or targets that are burst-sensitive, while higher rates maintain robust throughput where bursts are less impactful.

### The role of periodic firing in the SUB in spatial cognition

Speed and direction of movement are critical for path integration, and both inputs have been proposed as essential for grid cells. This could be provided by GC ensembles (53). From a mechanistic perspective, the computation of speed may rely on theta oscillations, which are strongly correlated with velocity, as well as on speed cells (54–58). Moreover, the direction of movement is of great relevance (59). This directional input may be provided by axis cells described in the SUB (60). Complementary to these inputs, the speed and directional signals could also be shaped by periodic cells in the MEC, pre-parasubiculum and SUB units reported here (60,61). The redundancy of different speeds and direction inputs might provide a more robust spatial integration for the navigation across different conditions, for example across light-dark transitions.

Our new evidence of spatially periodic cells suggests that the SUB may contribute to generating spatial representations by supporting periodic cells and GC in the pre-parasubiculum and MEC. This is supported by anatomical data that confirms the strong afferences sent by the SUB to the pre- and parasubiculum and the deep layers of the MEC (16,62). Additionally, the SUB might provide input from boundary vector cells (BVC) and periodic neurons to hippocampal place cells through its back projection to the CA1 area (63). In this way, circuits involving SUB-MEC, SUB-CA1, might support the transformation of the grid produced by alterations of the geometry of space (64). Finally, SUB spatial representation appears to depend on thalamic inputs and might meditate changes in GC due to thalamic lesions also known to be relevant for GC in MEC (65).

In addition, two interneurons also met our grid/periodicity criteria, indicating that inhibition can participate directly in spatially structured coding. This is consistent with recent publication (25) demonstrating a cooperative role for interneurons in shaping spatial representations.

### Data limitation and context of the results

One limitation of the tetrode cohort was the small N, with grid-like firing observed in two mice. Nevertheless, those units were stable across sessions and met stringent isolation criteria (refractory-period violations and cluster-overlap probability). In one implant, histology was incomplete, and the tetrode tip was not unambiguously visualized; depth was inferred from robust ripple activity, raising the possibility of partial sampling of adjacent regions (e.g., presubiculum), where ripples also occur (66). However, we obtained only one neurons from this implant. To address these concerns, we added an independent multisite probe cohort that provided higher yield and proximodistal coverage. In this dataset, 7/67 spatially modulated units (10.4%) met our priori periodic/grid criteria (5 pyramidal, 2 interneurons), an incidence that closely matched the tetrode dataset (4/28 = 14.3%) under identical analysis and shuffle thresholds. The replication across technologies and animals, together with consistent quality criteria and cell-type classification, substantially mitigates the small-sample caveat and supports the conclusion that periodic firing is a reproducible feature of mouse subiculum.

## Conclusion

Here we present evidence of spatial periodic firing in the SUB. However, their response to complex spatial manipulation or their relationship with BVC, GC and place need to be determined. The SUB is of great relevance for the integration of the spatial code, episodic memory and dimensions related to the ongoing tasks. Further work should aim to investigate the role of the SUB in memory consolidation and spatial coding and in other cognitive processes.

## Author Contributions

JRBM designed the experiments. JRBM and PA, GR, GC, MS developed experiments, and implemented the analysis toolbox. FJMP and ME contributed to the implementation of the analysis and discussed the results. AF, VB and LM supported the project providing resources for its development and discussed the results. JBRM wrote the paper, and the manuscript was further reviewed, and data discussed by JRBM, PA, AF, and RS.

## Acknowledgments

We would like to thank Liset Menendez de la Prida and Professor for her comments on this manuscript and their unconditional support for our work. Moreover, we want to thank Jim Donnett (Axona LTD) who was always available to help us with his technical support.

